# Physiological and Biochemical Responses of Grafted Tomato Plants to Salinity Stress: Evidence from the Syrian Coast

**DOI:** 10.64898/2026.06.02.729545

**Authors:** Najwa Ahmed, Ramzi Murshed, Safaa Najla

**Affiliations:** University of Damascus, Faculty of agriculture, Department of Horticulture, Damascus, Syria

**Keywords:** rootstock, tomato, grafting, osmotic adjustment, salt stress

## Abstract

The research was carried out in Tartus (Syria), in a plastic greenhouse during the season 2024-2025. Two tomato hybrids “Levovil and Shannon” were grafted onto two tomato “Spirit and Maxifort” rootstocks. Four levels of salinity (0, 50, 100 and 150 mM NaCl) were applied on the hybrids and grafted plants.

Salinty stress induced physiological and biochemical changes in plant. At salinity level of 150 mM, the absolute value of the osmotic pressure increased to -1.18 and -1.12 Mpa, in Levovil and Shannon hybrids, as compared to -0.80 and - 0.74 Mpa in the controls, respectively. While 150 mM salinity level led to an increase of the contents of dry matter, proline, sugars and chlorine, it caused a decrease of K: Na ratio in the two hybrids (1.99 and 1.8) as compared to the controls (5.29 and 4.68, respectively). The salinity stress at 150 mM, reduced the yield of the two hybrids (26.26 and 25.28 kg/m^2^) as compared to the controls (33.90 and 33.11 kg/m^2^, respectively).

The tomato scions “Levovil and Shannon hybrids” grafted onto the “Spirit and Maxifort” rootstocks enhanced the osmotic adjustment phenomenon, especially when the two hybrids were grafted onto Spirit. While tomato grafting had a significant effect on the accumulation of some osmotic compounds such as proline and sugars, in addition to increasing K: Na ratio, it reduced the absolute value of the osmotic pressure of the plant and its sodium content. Therefore, the yield increased in grafted tomato plants compared with non-grafted plants under saline conditions.

The results of PCA analysis showed that the first two principal components (PC1 and PC2) explained 89.08% of the total variation, wherease all the studied parameters were well represented. The dendrogram was obtained based on the same variables as PCA, and it led to classify “Spirit” rootstock as tolerant to salt stress, “Maxifort” rootstock as half tolerant, and the non-grafted hybrids “Levovil and Shannon” as sensitive to salt stress.

## Introduction

Tomato (*Solanum Lycopersicon* L.) is the second important vegetable in the world after potato, due to its nutritional and economic importance (FAO 2020). It is a model for physiological and genetic studies (Lim et al. 2016). Worldwide, the area planted with tomato reached to 5.1 million hectares, with a production of 186.8 million tons (FAO 2020).

In Syria, salinity poses a great danger as long as large areas are left out of agricultural investment, due to the absence of effective drainage systems and using water sources of poor quality and/or intertfering with sea water, as the case of several greenhouses along Syrian coast (Annual Agricultural Statistical Group 2018). According to USDA-ARS (2008), soils are cosidered as saline when the ECe is 4 dS/m or more (i. e. approximately 40mMNaCl). This level of salinity significantly reduces the yield of most crops.

Several studies have shown that salinity reduces plant growth by reducing leaf area, plant height, number of leaves, pore density and transpiration rate (Romero-Aranda et al. 2001). While dry matter accumulates in the plant (Gumi et al. 2013). The reduction of plant growth under salinity conditions occurs in two phases, the osmotic phase which starts immediately after the salt concentration in the soil increases to a threshold level, and the ion-specific phase which starts when salt accumulates to toxic concentrations in the old leaves (Hasegawa et al. 2000, Munns and Tester 2008). Furthermore, the salinity causes a disorder of ionic balance due to a decrease in the absorption rate of nutrient mineral elements (i.e. K^+^, Ca^++^, Mg^++^), which may be associated with a decrease in the transfer rate to plant branches (Serrano 1999). The vegetables differ in the tolerance to saltinity stress according to their ability to synthesize and accumulate the organic solutes (Gupta and Huang 2014). While some studies classify tomato as a salt moderately tolerant plant (Singh et al. 2012), other studies classify it as a salt-sensitive plant (Coban et al. 2020).

To solve salinity problem several techniques have been introduced such as, developing salinity-tolerant hybrids (Zheng et al. 2016). However, this technique achieved limited success due to the physiological and genetic complexity of salinity tolerance in plants (Lim et al. 2016). In this context, grafting on salt-tolerant rootstocks can be a procedure of the sustainable agriculture because it is safe and environmentally friendly (Kumar et al. 2017). Grafting increases growth rate (Balliu et al. 2007) and productivity of plant (Al-Harbi et al. 2017). In addition, it improves the morphological characteristics of the plants (Singh et al. 2020), and increases the rate of nutrients absorption, which is reflected in an increase in the rate of photosynthesis (Feng et al. 2019). On the other hand, grafting on salt tolerant rootstocks helps to alleviate negative impacts of salininty by increasing plant water content and improving fruits quality (Balliu et al. 2007; Colla et al. 2010). Several studies demonstrated that rootstocks with vigorous root systems produce more cytokinins which promote the growth of tomato plant under salinity conditions (Albacete et al. 2015; Albornoz et al. 2020). The ability of plant to control the absorption, transport, sequestration and accumulation of ions such as Na^+^, Cl^™^ and K^+^, is one of the most important mechanisms of salinity tolerance (Tester and Davenport 2003; Parida and Das 2005; Munns and Tester 2008; Jha et al. 2010). In addition, the phenomenon of osmotic adjustment, whereby the plant increases the production of compatible solutes such as amino acids and sugars (Almeida et al. 2014) is an effective mechanism to tolerate the plant to salinity. These compounds accumulate in the cytoplasm and contribute to maintain the turgor pressure and the structure of the cell (Colla et al. 2010; Conde et al. 2011).

Recently, tomato cultivation in the Syrian coastal region witnessed a noticeable decline in production, despite the farmers’ tendency to use grafted plants. However, they focus only on the diseases biological - resistant rootstocks, without taking into account the abiotic problems, which the region suffers from, such as salinity. This research aims to investigate the effect of tomato grafting on salinity tolerance by evaluating some compoenets envolvoled in the osmotic adjustment mechanism. Thus, this research can provide an information base for future studies on the possibility of adopting certain rootstocks of tomatoes in the salinized Syrian coastal area.

## Materials and Methods

### Experimental site and plant materials

The research was carried out in a plastic greenhouse in Tartus (Syria). The greenhouse was equipped with heating and ventilation systems to provide a mean daytime/night temperatures of 25 °C/15 °C and a relative humidity between 70 and 80%.

Two scions (Shannon F1 and Levovil F1) and two rootstocks (Spirit F1 and Maxifort F1) of tomato plant were used. The scions seeds were sown on 8/8/2021 and the rootstocks seeds were sown 3 days later, in trays (3 x 3 x 7 cm). A sterile turbe material was used as a culture medium. The growth medium was irrigated with water containing soluble fertilizer (20:20:20%, N: P: K) at a rate of 1 g/L. When the seedlings reached the appropriate size (2-3 true leaves), the tube grafting was applied on the healthy seedlings of similar diameter. After grafting, the trays were transferred to the healing chamber (temperature 24°C, relative humidity 90%), for 5 days. After healing, the temperature was increased and the relative humidity was gradually decreased for 4 days for the purpose of acclimatization of seedlings to the external conditions. Afterwards, the seedlings were planted in the greenhouse on 9/30/2024. The planting distance between rows was 70 cm, and between plants in the row was 40 cm. The agricultural density was 3.5 plants/m^2^. The seedlings were watered immediately and the common cultivation practices were applied.

Four levels of salinity (0, 50, 100 and 150 mM of pure NaCl) were applied to four grafting combinations (Shannon and Levovil/Spirit and Maxifort), after the appearance of the first flower cluster.

At the fruiting stage, 5 representative plant samples were randomly chosen from each replicate in order to achieve the measrements. Plant osmotic pressure (MPa) was carried out by the osmometer (OM 815, Vogel GmbH and Co. KG, Germany). For determining the dry matter content, the leaves were oven dried at 60ºC for 48 hrs and weighed according to Murshed et al. (2015). Na^+^ and K^+^ (ppm) were estimated by the flame spectrometer according to Arbaoui et al. (2020) and Cl^-^ (ppm) was determined according to the Gaines et al. (1984). Proline content in leaves (mg/g FW) was estimated using a spectrophotometer at wavelength 528 nm (Dreier and Goring 1974). Sugars content of leaf were estimated by enzymatic methods according to Mannheim (1989). The yield (kg/m^2^) was estimated per square meter.

### Experiment Design and Statistical Analysis

The experiment was designed according to completely randomized sectors. It contained 4 grafting combinations and two non-grafted hybrids (6 status of the plant) and 4 levels of salinity. Each treatment contained 3 replicates and each replicate consisted of 16 plants, so the total number of plants was 1152. The data were analyzed using R-Project program, version 4.0.3 and Fisher’s test was used to calculate the value of the least significant difference (LSD) between the variables at 95% confidence level.

To evaluate relationships between the factors and to detect the most important factors influencing tomato response, PCAs were run. To determine the tolerance of grafted and non-grafted hybrids to salt stress, cluster analysis was performed according to Murshed et al. (2015):

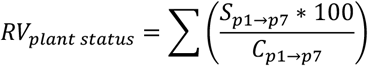

Where, *RV*_*plant status*_ is the sum of the relative values of the plant status (grafted or non-grafted hybrid), *S*_*p*1→*p*7_ is the value of the studied parameter (7 parametres) in the stressed plant, *C*_*p*1→*p*7_ is the value of the studied parameter in plant control.

## Results

### 1. Effect of salt stress and grafting on plant osmotic pressure (π, MPa)

Table 1 showed that salt stress (150, 100 and 50 mM) in Levovil F1, led to a significant increase in the absolute value of π (-1.18, -1.02 and -0.94 MPa, respectively) as compared to the control (-0.80 MPa). Grafting on both “Spirit and Maxifort” rootstocks lowered the osmotic pressure (-0.94 and -0.93 MPa, respectively) as compared to non-grafted plants (MPa -1.09). Regarding the effect of the interaction (stress X grafting), it is inferd that grafting can reduce significantly the negative effects of salinity up to 100 mM, while tomato grafting at 150 mM did not enhance salt tolerance as compared to the non-grafted plants at the same stress level (-1.21 MPa).

**Table 1:**
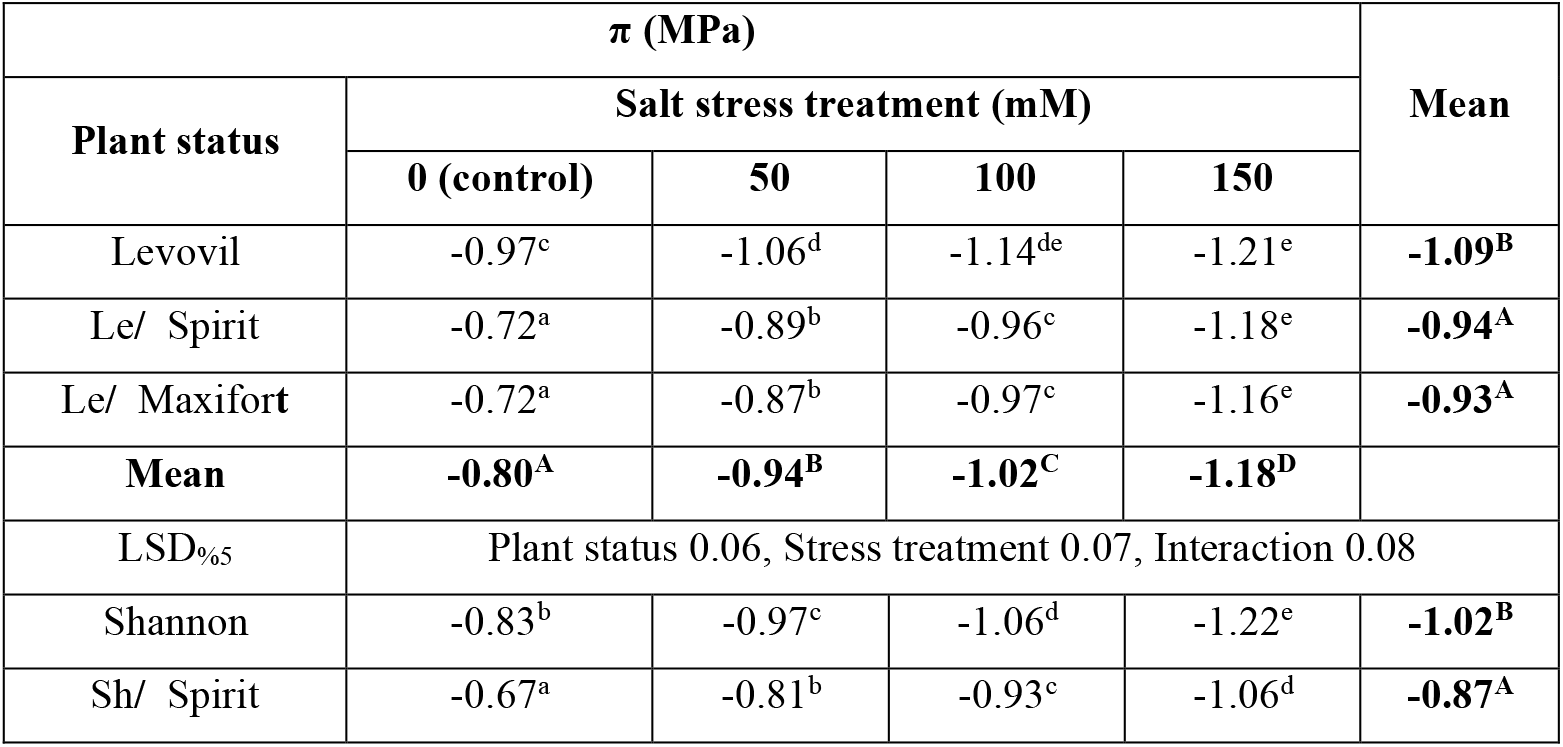
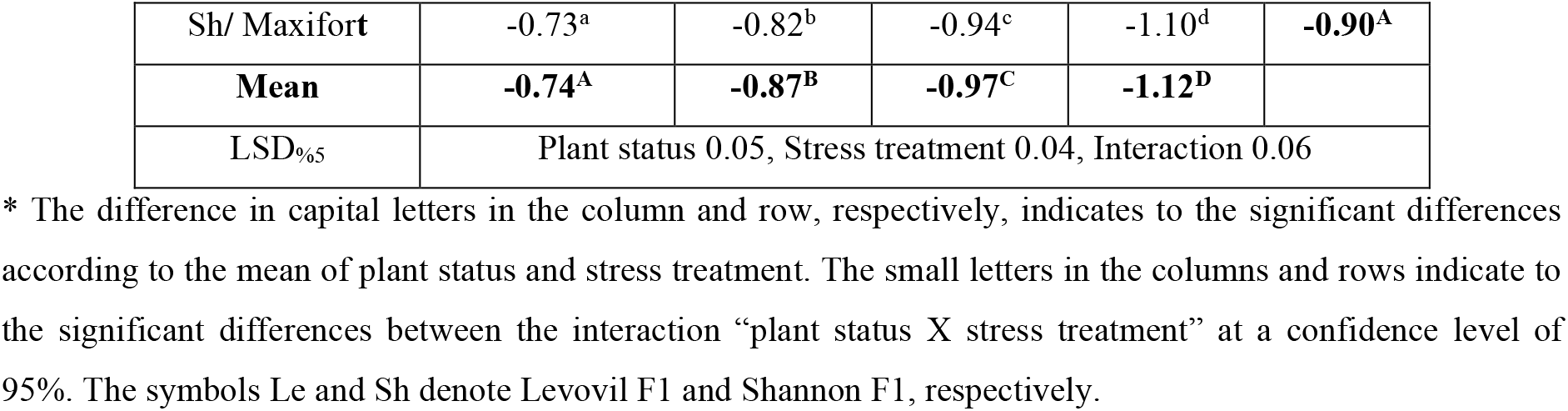
Effect of salt stress and grafting on plant osmotic pressure (MPa).

The same trend of salt stress effect was recorded in Shannon F1 (table 1). π increased gradually and significantly with stress increase (-0.87, -0.97 and -1.12 MPa, respectively) as compared to the control (0.74 MPa). Also, grafting onto “Spirit and Maxifort” rootstocks reduced π (-0.87 and -0.90 MPa, respectively) as compared to non-grafted plants (MPa -1.02). Regarding the interaction, it is noted that grafting at all stress levels decreased significantly the osmotic pressure as compared to the non-grafted plants.

### 2. Effect of salt stress and grafting on plant dry matter content (%)

Table 2 shows that salt stress in Levovil hybrid did not affect the dry matter content significantly, except at a severe stress level (150 mM), where it reached to 10.91% as compared to the control (9.66%). Grafting onto “Spirit and Maxifort” rootstocks increased significantly the dry matter content by 1.3 and 1.2 times, respectively, as compared to the non-grafted Levovil (8.99%). Grafting on the two rootstocks at 50 and 100mM showed a significant increase in dry matter content, while at 150 mM no significant effect was observed comparing to non-grafted Levovil at the same salinity levels.

**Table 2:**
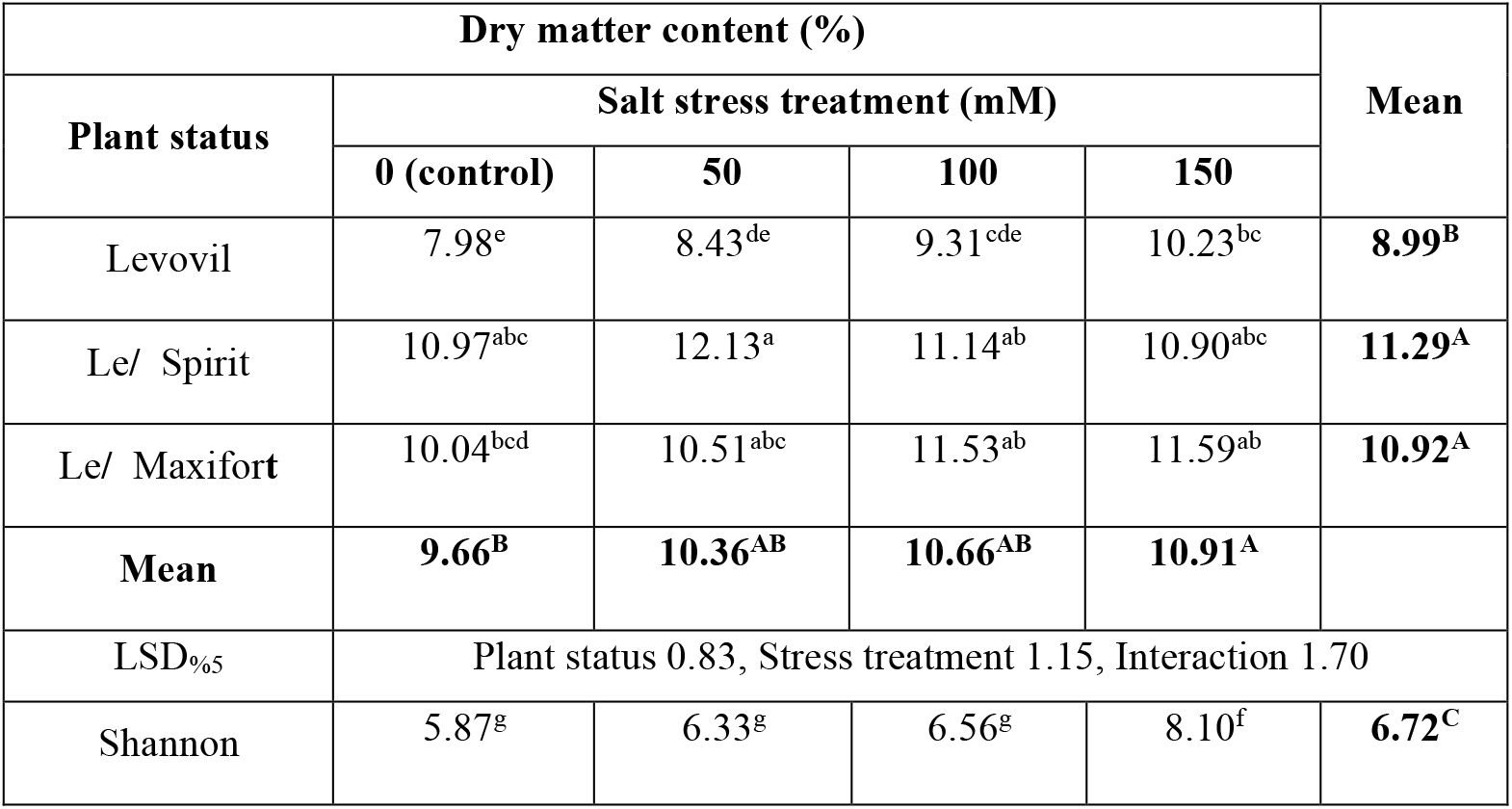
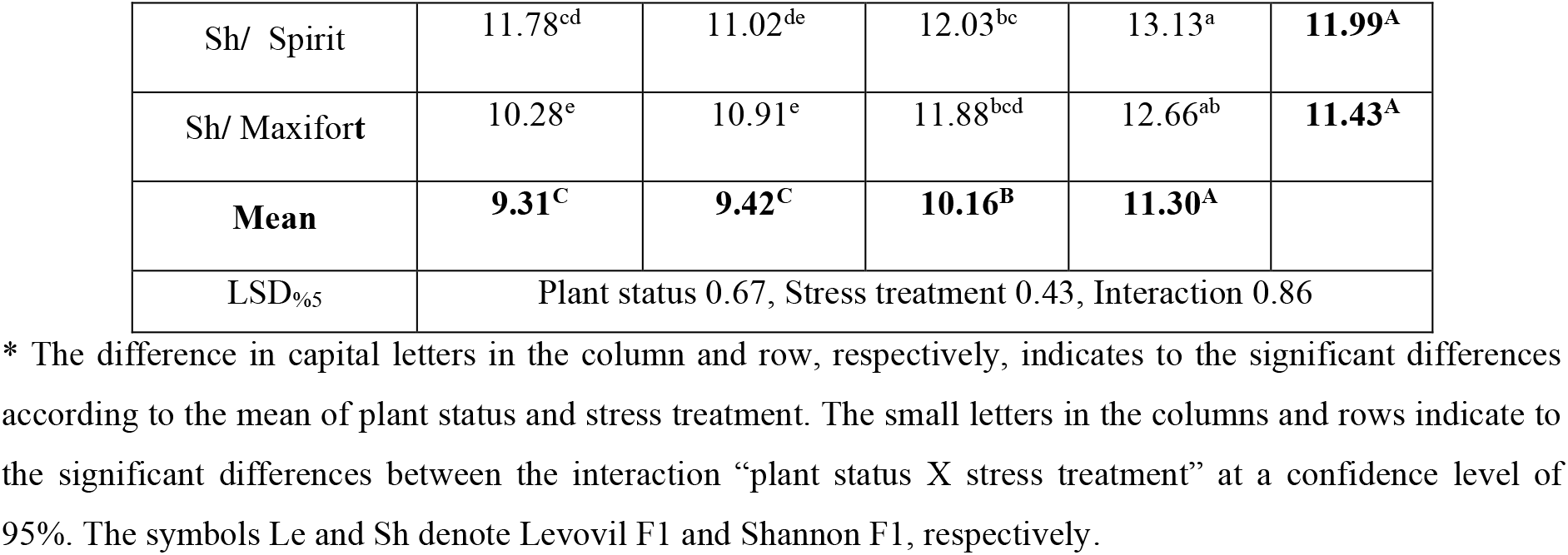
Effect of salt stress and grafting on plant dry matter content (%):

In Shannon hybrid, the salt stress at 100 and 150 mM (table 2), led to a significant increase in the dry matter content by 1.1 and 1.2 times as compared to the control (9.31%). Grafted Shannon onto “Spirit and Maxifort” contributed to an increase in the studied parameter (11.99 and 11.43%, respectively) as compared to the non-grafted Shannon (6.72%). The grafted Shannon, on both rootatocks and under all salinity levels, showed a significant increase in the dry matter content as compared to non-grafted scion at the same salinity levels.

### 3. Effect of salt stress and grafting on the ratio of K: Na in leaves

The salt stress, only at 100 and 150 mM, reduced significantly the ratio of K: Na of Levovil leaves (table 3) by 35.64 and 40.69 %, as compared to the control (3.76). Whereas, grafting only on “Spirit” rootstock increased significantly K: Na ratio by 1.19 times as compared to the non-grafted Levovil (2.67). For the interaction, grafting on the two rootstocks show a positive result in protecting the plant from salt stress only at 100 mM, where a significant increase in the studied indicator was observed as compared to the non-grafted scion at the same salinity level.

**Table 3:**
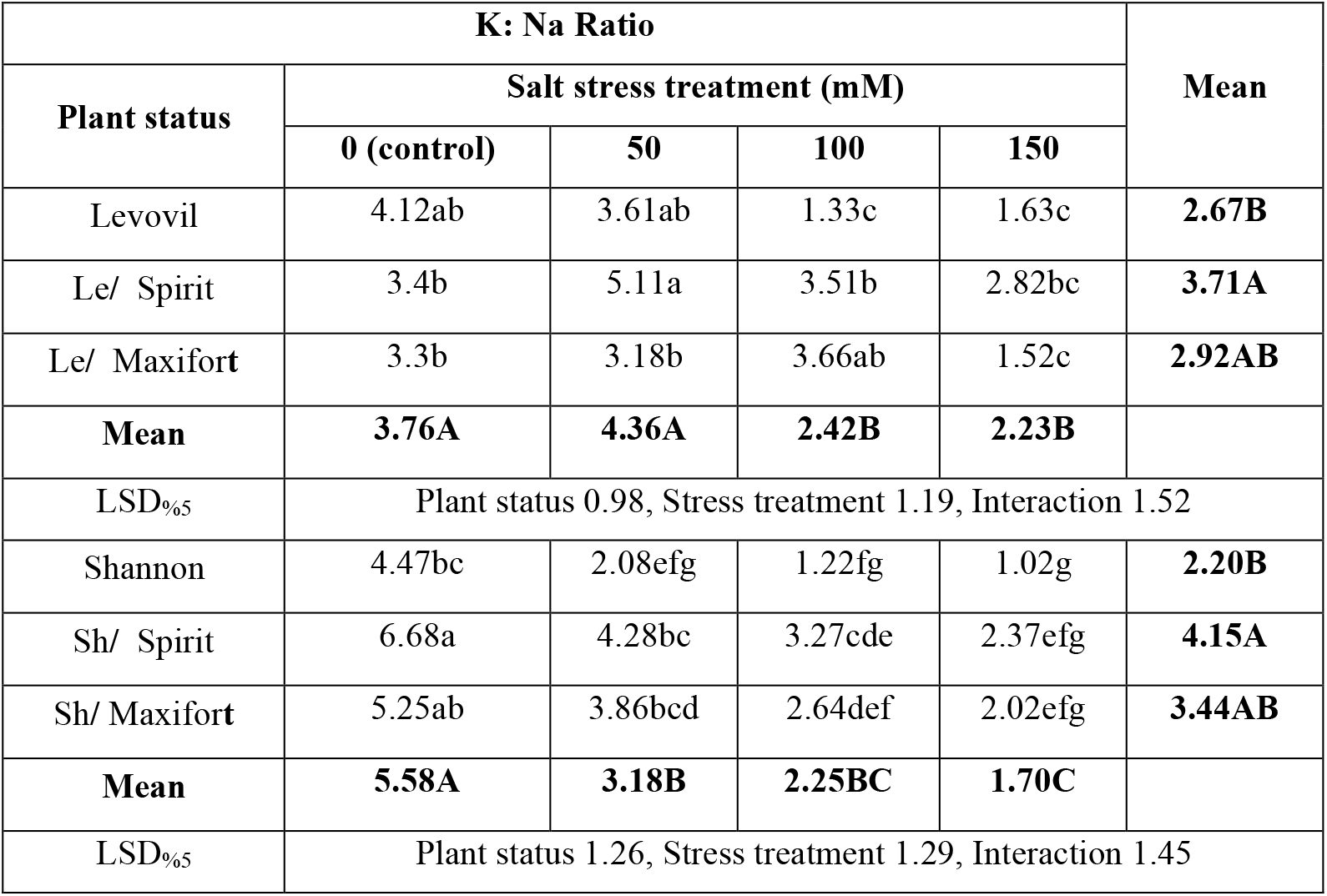
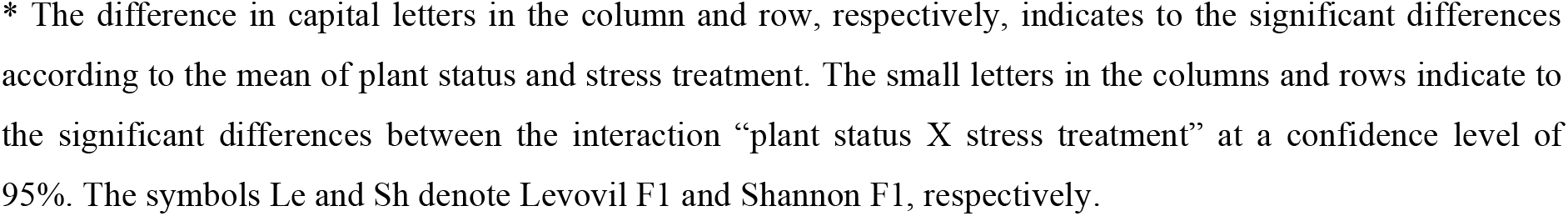
Effect of salt stress and grafting on the ratio of K: Na in leaves.

| K: Na Ratio |  |  |  |  | Mean |
| --- | --- | --- | --- | --- | --- |
| Plant status | Salt stress treatment (mM) |  |  |  |  |
|  | 0 (control) | 50 | 100 | 150 |  |
| Levovil | 4.12ab | 3.61ab | 1.33c | 1.63c | 2.67B |
| Le/ Spirit | 3.4b | 5.11a | 3.51b | 2.82bc | 3.71A |
| Le/ Maxifort | 3.3b | 3.18b | 3.66ab | 1.52c | 2.92AB |
| Mean | 3.76A | 4.36A | 2.42B | 2.23B |  |
| LSD<sub>5</sub> | Plant status 0.98, Stress treatment 1.19, Interaction 1.52 |  |  |  |  |
| Shannon | 4.47bc | 2.08efg | 1.22fg | 1.02g | 2.20B |
| Sh/ Spirit | 6.68a | 4.28bc | 3.27cde | 2.37efg | 4.15A |
| Sh/ Maxifort | 5.25ab | 3.86bcd | 2.64def | 2.02efg | 3.44AB |
| Mean | 5.58A | 3.18B | 2.25BC | 1.70C |  |
| LSD<sub>5</sub> | Plant status 1.26, Stress treatment 1.29, Interaction 1.45 |  |  |  |  |

\* The difference in capital letters in the column and row, respectively, indicates to the significant differences according to the mean of plant status and stress treatment. The small letters in the columns and rows indicate to the significant differences between the interaction “plant status X stress treatment” at a confidence level of 95%. The symbols Le and Sh denote Levovil F1 and Shannon F1, respectively.

### 4. Effect of salt stress and grafting on Cl^-^ content (%) of leaves

Table 4 shows that the salt stress at 100 and 150 mM increased signifcantly the leaf Cl^-^ content of Levovil F1 by 1.3 and 1.2 times, respectively, as compared to the control (0.58%). However, grafting only on Maxifort reduced signifcantly the leaf Cl^-^ content (0.63%) as compared to the non-grafted scion (0.71%). Only the combination Le**/** Maxifor**t** recorded a significant reduction in the studied parameter at 50 mM as compared to the non-grafted scion at the same salinity level.

**Table 4:** Effect of salt stress and grafting on the Cl^-^ content (%) of leaves.

| Cl<sup>-</sup> (%) |  |  |  |  | Mean |
| --- | --- | --- | --- | --- | --- |
| Plant status | Salt stress treatment (mM) |  |  |  |  |
|  | 0 (control) | 50 | 100 | 150 |  |
| Levovil | 0.63cde | 0.66bcd | 0.74ab | 0.81a | **0.71A** |
| Le/ Spirit | 0.56de | 0.59de | 0.64bcde | 0.77a | **0.64A** |
| Le/ Maxifort | 0.55e | 0.54e | 0.71abc | 0.72abc | **0.63B** |
| Mean | **0.58B** | **0.6B** | **0.7A** | **0.77A** |  |
| LSD<sub>%5</sub> | Plant status 0.07, Stress treatment 0.08, Interaction 0.10 |  |  |  |  |
| Shannon | 0.57cde | 0.66abcd | 0.72ab | 0.77a | **0.68A** |
| Sh/ Spirit | 0.56de | 0.53de | 0.63bcde | 0.74ab | **0.62A** |
| Sh/ Maxifort | 0.54de | 0.51e | 0.56de | 0.70abc | **0.58A** |
| Mean | **0.56B** | **0.57B** | **0.64AB** | **0.74A** |  |
| LSD<sub>%5</sub> | Plant status 0.10, Stress treatment 0.11, Interaction 0.13 |  |  |  |  |

\* The difference in capital letters in the column and row, respectively, indicates to the significant differences according to the mean of plant status and stress treatment. The small letters in the columns and rows indicate to the significant differences between the interaction “plant status X stress treatment” at a confidence level of 95%. The symbols Le and Sh denote Levovil F1 and Shannon F1, respectively.

The salt stress caused, only at 150 mM, a significant increase in Cl^-^ content of Shannon leaves (0.74%) as compared to the control (0.56%). While grafting Shannon on both rootstocks did not affect the studied parameter (0.62 and 0.58% in Sh**/**Spirit and Sh**/**Maxifor**t**, respectievly) as compared to non-grafted Shannon (0.68%). In term of interaction, grafting, only on Maxifort rootstocks, reduced the negative effect of salt stress at 50 and 100 mM by decreasing Cl^-^ content in comparison with the non-grafted Shannon at the same salinity level (table 4).

### 5. Effect of salt stress and grafting on leaf proline content (mg/g Fw)

The salt stress (100 and 150 mM) increased significantly the proline content of Levovil by 1.4 and 1.7 times, respectively (table 5), as compared to the control (0.97 mg/g Fw). Also, grafting onto Spirit and Maxifort led to a significant increase in the studied parameter by 1.4 and 1.5 times, respectively, as compared to the non-grafted Levovil (0.98 mg/g Fw). Regarding the interaction, the combination Le/Maxifort mitigated significantly the effect of salt stress at all levels by increasing the proline content, while the combination Le/Spirit mitigated significantly the effect of salt stress only at 50 and 150 mM as compared to the non-grafted scion at the same salinity level.

**Table 5:**
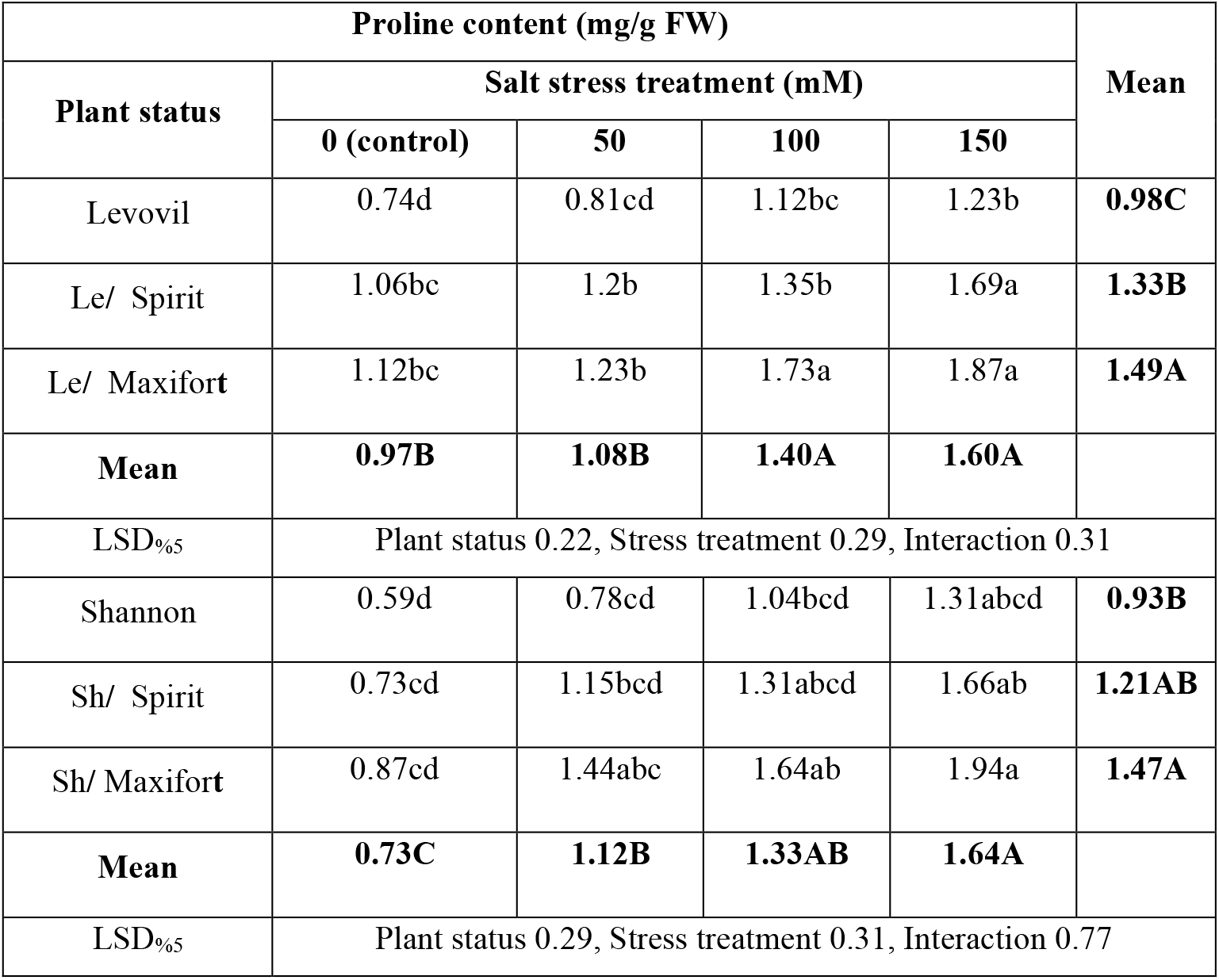
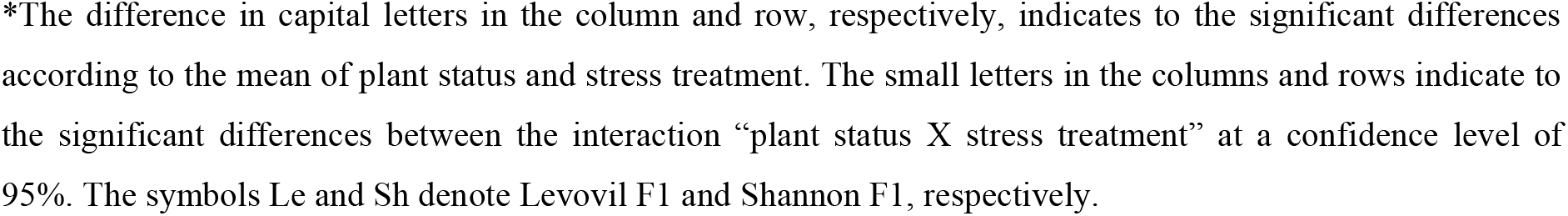
Effect of salt stress and grafting on plant proline content (mg/g FW)

| Proline content (mg/g FW) |  |  |  |  | Mean |
| --- | --- | --- | --- | --- | --- |
| Plant status | Salt stress treatment (mM) |  |  |  |  |
|  | 0 (control) | 50 | 100 | 150 |  |
| Levovil | 0.74d | 0.81cd | 1.12bc | 1.23b | 0.98C |
| Le/ Spirit | 1.06bc | 1.2b | 1.35b | 1.69a | 1.33B |
| Le/ Maxifort | 1.12bc | 1.23b | 1.73a | 1.87a | 1.49A |
| Mean | 0.97B | 1.08B | 1.40A | 1.60A |  |
| LSD<sub>%5</sub> | Plant status 0.22, Stress treatment 0.29, Interaction 0.31 |  |  |  |  |
| Shannon | 0.59d | 0.78cd | 1.04bcd | 1.31abcd | 0.93B |
| Sh/ Spirit | 0.73cd | 1.15bcd | 1.31abcd | 1.66ab | 1.21AB |
| Sh/ Maxifort | 0.87cd | 1.44abc | 1.64ab | 1.94a | 1.47A |
| Mean | 0.73C | 1.12B | 1.33AB | 1.64A |  |
| LSD<sub>%5</sub> | Plant status 0.29, Stress treatment 0.31, Interaction 0.77 |  |  |  |  |

\*The difference in capital letters in the column and row, respectively, indicates to the significant differences according to the mean of plant status and stress treatment. The small letters in the columns and rows indicate to the significant differences between the interaction “plant status X stress treatment” at a confidence level of 95%. The symbols Le and Sh denote Levovil F1 and Shannon F1, respectively.

Table 5 shows, that the higher the stress, the higher the proline content in Shannon leaves (1.5, 1.8 and 2.2 times, respectively) as compared to the control (0.73 mg/g Fw). As well as grafting only on “Maxifort” rootstock led to a significant increase in the studied parameter by 1.6 times as compared to the non-grafted hybrid (0.93 mg/g Fw). Regarding the interaction, none of the combinations had recorded any differnece as compared to non-grafted scion at the same salinity level.

### 6. Effect of salt stress and grafting on plant sugars content (%)

The higher the salinity stress, the higher the sugars content in Levovil plant as compared to the control (0.46 %). Grafting on “Spirit and Maxifort” resulted in a significant increase of sugars content (.058 and 0.56%) as compared to the non-grafted Levovil (0.52%). Table 6 shows that both combinations did not record any effects under all stress levels, as compared to non-grafted scion at identical salinity levels.

**Table 6:** Effect of salt stress and grafting on plant sugars content (%):

| Sugars content |  |  |  |  | Mean |
| --- | --- | --- | --- | --- | --- |
| Plant status | Salt stress treatment (mM) |  |  |  |  |
|  | 0 (control) | 50 | 100 | 150 |  |
| Levovil | 0.40e | 0.47d | 0.58c | 0.64ab | **0.52B** |
| Le/ Spirit | 0.51d | 0.52d | 0.60bc | 0.68a | **0.58A** |
| Le/ Maxifort | 0.48d | 0.50d | 0.61bc | 0.65ab | **0.56A** |
| Mean | **0.46D** | **0.50C** | **0.60B** | **0.66A** |  |
| LSD%<sub>5</sub> | Plant status 0.03, Stress treatment 0.02, Interaction 0.05 |  |  |  |  |
| Shannon | 0.43d | 0.45d | 0.51c | 0.60b | **0.50C** |
| Sh/ Spirit | 0.48cd | 0.51c | 0.61ab | 0.66a | **0.57A** |
| Sh/ Maxifort | 0.46d | 0.48cd | 0.58b | 0.65a | **0.54B** |
| **Mean** | **0.46C** | **0.48C** | **0.57B** | **0.64A** |  |
| LSD<sub>5%</sub> | Plant status 0.02, Stress treatment 0.03, Interaction 0.04 |  |  |  |  |

\* The difference in capital letters in the column and row, respectively, indicates to the significant differences according to the mean of plant status and stress treatment. The small letters in the columns and rows indicate to the significant differences between the interaction “plant status X stress treatment” at a confidence level of 95%. The symbols Le and Sh denote Levovil F1 and Shannon F1, respectively.

The salt stress at 100 and 150 mM (table 6) increased significantly the sugars content of Shannon (0.57 and 0.64%, respectively) as compared to the control (0.46 %). Grafting on both “Spirit and Maxifort” rootstoks increased significantly the studied parameter (0.57 and .054%, respectively), as compared to the non-grafted hybrid (0.50 %). The combination Sh**/** Spirit contributed to mitigating the effect of salt stress at all levels, while the combination Sh**/** Maxifort contributed to mitigating the effect of stress at 100 and 150mM, as compared to non-grafted plants at identical salinity levels.

### 7. Effect of salt stress and grafting on the yield (kg/m^2^)

Table 7 shows that salinity stress (50, 100 and 150 mM) decreased significantly the yield of Levovil by 1.1, 1.2 and 1.3 times as compared to the control (33.90 kg/m^2^). However, grafting onto “Spirit and Maxifort” rootstocks increased the yield by 1.4 and 1.2 times, respectively, as compared to the non-grafted hybrid (24.83 kg/m^2^). Noting that grafting on Spirit recorded significant differences as compared to grafting on Maxifort. Morover, both combinations (Le/Spirit and Le/Maxifort) reduced the effect of all levels of salt stress, as compared to non-grafted plants at identical salinity levels.

**Table 7:** Effect of salt stress and grafting on the yield (kg/m^2^)

| Yiled (kg/m<sup>2</sup>) |  |  |  |  | Mean |
| --- | --- | --- | --- | --- | --- |
| Plant status | Salt stress treatment (mM) |  |  |  |  |
|  | 0 (control) | 50 | 100 | 150 |  |
| Levovil | 28.13de | 26.84e | 22.76f | 21.57f | 24.83C |
| Le/ Spirit | 38.55a | 35.19ab | 31.37c | 29.83cd | 33.74A |
| Le/ Maxifort | 35.03b | 32.34c | 28.77d | 27.39de | 30.88B |
| Mean | 33.90A | 31.46B | 27.63C | 26.26C |  |
| LSD<sub>%5</sub> | Plant status 2.53, Stress treatment 2.34, Interaction 2.88 |  |  |  |  |
| Shannon | 29.27c | 27.26c | 23.96d | 21.61d | 25.53B |
| Sh/ Spirit | 34.92a | 31.98b | 28.41c | 27.37c | 30.67A |
| Sh/ Maxifort | 35.15a | 31.95b | 27.79c | 26.87c | 30.44A |
| **Mean** | **33.11A** | **30.4A** | **26.72B** | **25.28B** |  |
| LSD<sub>5%</sub> | Plant status 2.12, Stress treatment 2.23, Interaction 2.55 |  |  |  |  |

\* The difference in capital letters in the column and row, respectively, indicates to the significant differences according to the mean of plant status and stress treatment. The small letters in the columns and rows indicate to the significant differences between the interaction “plant status X stress treatment” at a confidence level of 95%. The symbols Le and Sh denote Levovil F1 and Shannon F1, respectively.

Likewise, it was observed that all levels (50, 100 and 150 mM) of saline stress reduced significantly the yield of Shannon (30.4, 26.72 and 25.28 kg/m^2^, respectively) as compared to the control (33.11 kg/m^2^). Grafting on Spirit and Maxifort increased significantly the yield by 1.20 and 1.19 times, respectively, as compared to the non-grafted scion (25.53 kg/m^2^), without recording significant differences between the two rootstocks. Both combinations (Le/Spirit and Le/Maxifort) contributed to mitigating the effect of salt stress at all levels, as compared to non-grafted plants at identical salinity levels.

### 8. Principal Component Analysis (PCA) and Agglomerative hierarchical clustering (AHC) of the measurements relative values

According to PCA (table 8), all the total varitaion has been dervied from 7 pricipal component axis. The first (PC1=59.81%) and second principal component (PC2=29.26 %) explained 89.08% of the total variation, while the rest of principle components (PC3, PC4, PC5, PC6 and PC7) had 10.92% of the total variation. PC1 and PC2 are more informative than the original variable, since its eigenvalue >1.

**Table 8:** Eigenvalues, variability and cumulative values.

|  | PC1 | PC2 | PC3 | PC4 | PC5 | PC6 | PC7 |
| --- | --- | --- | --- | --- | --- | --- | --- |
| Eigenvalue | 4.187 | 2.048 | 0.372 | 0.176 | 0.111 | 0.064 | 0.041 |
| Variability (%) | 59.815 | 29.263 | 5.316 | 2.518 | 1.592 | 0.917 | 0.580 |
| Cumulative (%) | 59.815 | 89.078 | 94.394 | 96.912 | 98.504 | 99.420 | 100.000 |

The quality of representation (cos^2^) values of plant measurements showed that the seven studied parameters (Os=0.894, DM=0.929, K:Na=0.719, Cl=0.819, Proline=0.476, Sugars=0.671, Yield=0.657) are well represented in PC1 and PC2 (Fig 1). The examination of Fig. 1 showed there is a positive relationship between the narrow-angle features (for example; Os with Cl and sugars, DM with prolin and sugars…etc), right angle features are not related to each other (for example; Os, Cl, K:Na and yield with DM…etc), and wide-angle features have negative relationships with each other (for example; Os with K:Na and yield, K:Na with Cl, Cl with yield).

**Fig 1.**
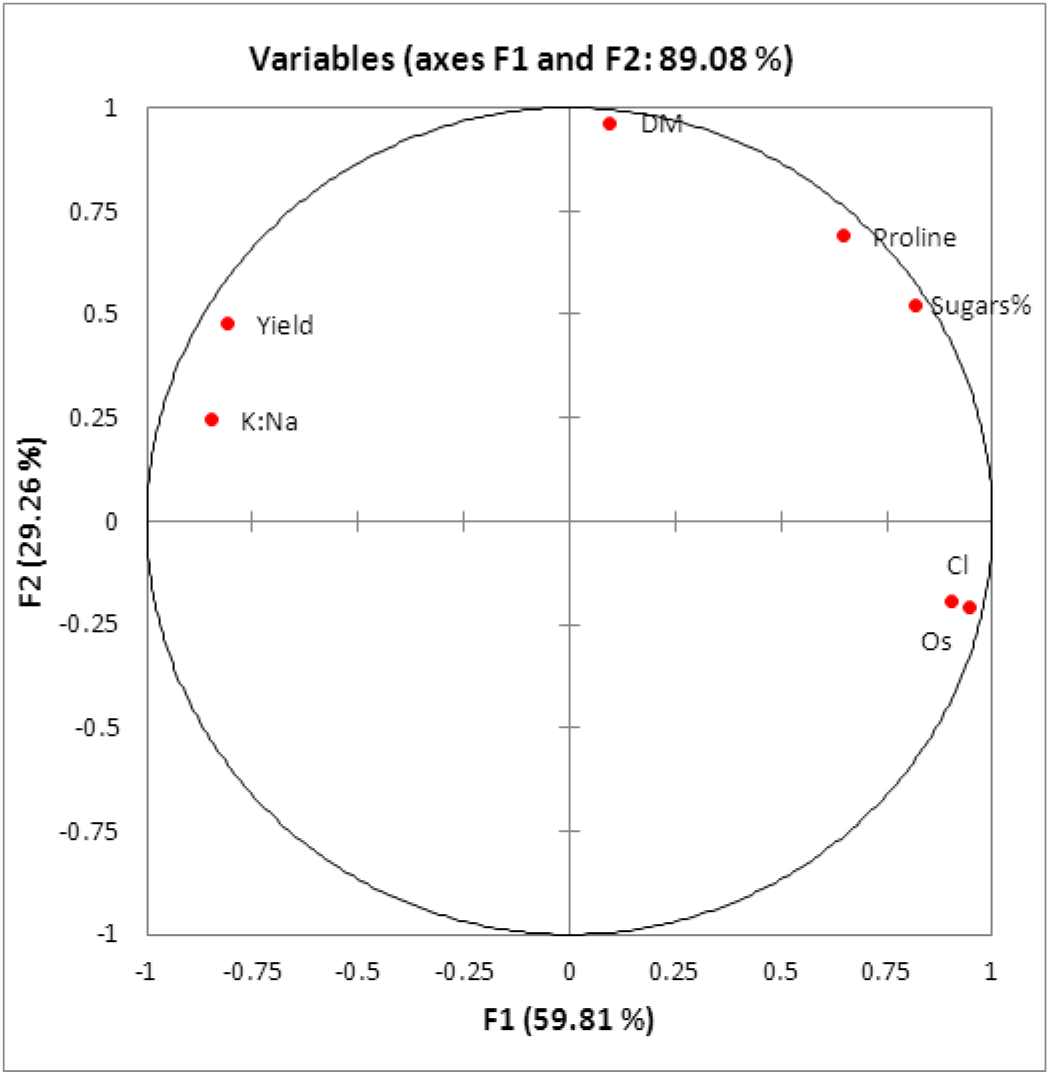
Quality of representation (cos^2^) correlation circle of plant parameters to PC1 and PC2. The plant parameters are osmotic pressure (Os), Dry matter content (DM), ratio K:Na, cloride content (Cl), proline content, sugar content and yield

The hierarchical cluster analysis, depending on the sum of the relative values of the seven studied criteria, classified the 6 plant status according to salt stress tolerance in three groups (Fig. 2): (i) salt tolerant group consisting of two combinations (Le/Spirit and Sh/Spirit); (ii) a moderately salt tolerant group including two combinations (Le/Maxifort and Sh/Maxifort); (iii) a salt susceptible group consisting of non-grafted Levovil and Shannon scions.

**Fig 2.**
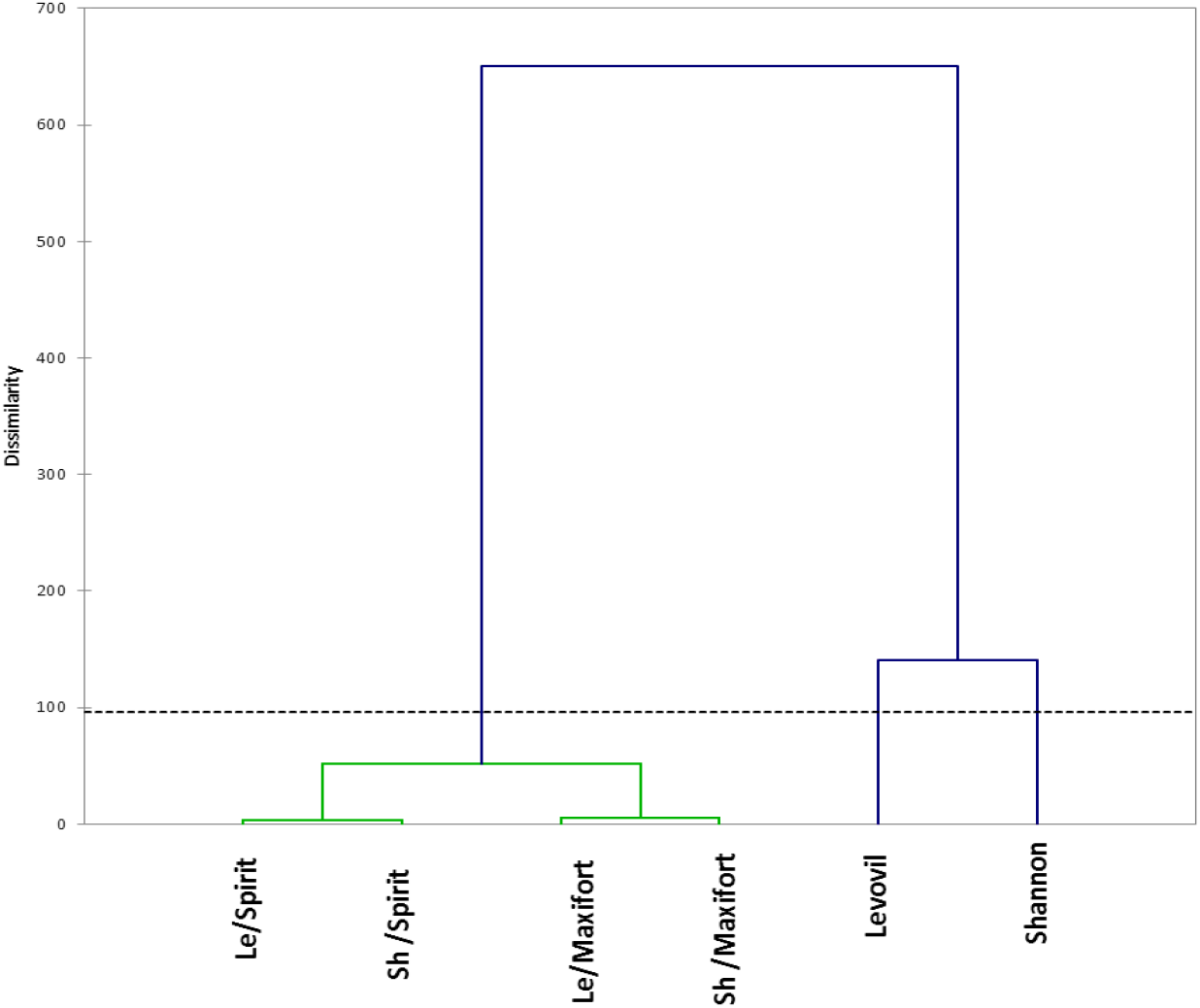
Cluster analysis of non-grafted and grafted tomato hybrids, according to their tolerance to salt stress based on the sum of the relative values of all studied parameters

## Disscuion

The salt stress affected the physiological and biochemical processes, and this can be explained by the plant water status which is considered as one of the main indicators in determining the plant response to stress (Munns et al. 2000; Ladewig et al. 2021). As a result of high concentration of soluble salts in the soil, the water potential gradient between the soil and the root system cells decreases, and the amount of water absorbed becomes insufficient to compensate the water lost by transpiration, and consequently plant osmotic pressure increases (table 1). On the other hand, Salinity led to an increase in plant contents of Na^+^ and Cl^-^ (Table 3 and 4); which the studies have proven their toxic effect by inhibiting the activity of many enzymes and hormones such as cytokines (Babu et al. 2012). Previous studies have shown that there is a competition between Na^+^ and K^+^ ions in terms of the ability to bind to special sites (Bhandal and Malik 1988).

The plant growth reduces as a result of decreasing the turgor pressure responsible for the cells elongation (Hasegawa et al. 2000). However, in plant tolerant to salt stress, the cell turgor is maintained by the phenomenon of osmotic adjustment, in which some osmotic and organic compounds such as potassium (table 3), proline (table 5), sugars (table 6), glycine betaine, proteins and polyamines…etc (Alarcon et al. 1994; Cola et al. 2010) are accumulated. In addition to their role in maintaining cell turgor, these compounds contribute to scavenging free radicals from the cell and maintain the stability of membrane proteins (Hou et al. 2021).

The decrease in yield, especially at 100 and 150 mM, can be explained by the decrease in growth indicators which leads to a decrease in photosynthesis products, and consequently the yield. In addition, the decrease in K^+^ content of the leaf (table 3) affects the number and weight of fruit, and consequently decreases the yield (Coban et al. 2020). It should be noted that salinity at 50mM did not significantly affect the yield of non-grafted hybrids (Levovil and Shannon) as compared to the controls, and this corresponds to many studies that advised that the salinity of medium should not exceed 50mM when growing tomatoes (Rodriguez-Ortega et al. 2019; Ladewig et al. 2021).

Grafting on both rootstocks increased signifcantly the yield as compared to non-grafted plants (24.83 and 25.53 kg/m^2^, respectively). Such an increase can be attributed to the higher rate of water and nutrients absorption by the vigorous rootstock (Ruiz et al. 1997; Estañ et al. 2009) as well as the plant hormones content, especially the cytokinins synthesized in the root system (Sharma and Zheng 2019). On the other hand, low osmotic pressure, Na^+^ and Cl^-^ contents (table 1, 3 and 4) and high sugars and proline contents (table 5 and 6) indicate the enhancement of the osmotic adjustment phenomenon and therefore an increase in yeild (Alarcon et al. 1994). The studies have confirmed that the lower Na^+^ and Cl^-^ content and the higher K^+^ content in the plant, the more tolerant the plant is to salt stress (Munns and Tester 2008; Huang et al. 2011). Therefore, tomato grafting on Spirit and Maxifort rootstocks can protect the plant from the negative effects of salinity, due to thier ability to reduce the transfer of the electrolytes (Na^+^ and Cl^-^) through the roots of the rootstocks to the aerial parts (Al-Harbi et al. 2017). Moreover, this result is consistent with the results of cluster analysis (Fig. 1) which showed that “Spirit” rootstock is a salt-tolerant plant and “Maxifort” rootstock is a moderately salt tolerant plant. However, the inconsistency of some results with previous studies can be explained by the difference in the compatibility between the rootstocks and the scions, as well as the difference in the experiment treatments and conditions.

The progress in breeding programs related to the salt stress tolerance of tomato still slow, since it is genetically complex trait (QTLs) (Foolad et al. 2001; Wang et al. 2021). Therefore, several studies have determined the tolerance of tomato varities to stress salt, using PCA and cluster analysis based on morphological, physiological or biochemical indicators (Dasgan et al. 2002; Seth 2018), and then compared with molecular assessment if possible.

In conclusion, in the Syrian coast, non-grafted hybrids could be used in saline conditions which do not exceed 50mM. However, at high salinity levels (100 and 150 mM), grafting on vigorous and salt -tolerant rootstocks is an urgant necessity. Spirit rootstock can be recommended, especially in conditions of high salinity, for its role in stimulating plant growth and increasing yield.

## Internet sites

Annual Agricultural Statistical Group, Ministry of Agriculture and Agrarian Reform in Syria (2018). http://moaar.gov.sy/main/archives/category.

FAO (2020). Food and Agriculture Organization of The United Nations. Information and Statistics Service.

Rome, FAO. http://faostat.fao.org.

The R Project for Statistical Computing. https://www.r-project.org

USDA-ARS. 2008. Research Databases. Bibliography on Salt Tolerance. George E. Brown, Jr. Salinity Lab. US Dep. Agric., Agric. Res. Serv. Riverside, CA. http://www.ars.usda.gov/Services/docs.htm?docid=8908.

